# SARS-CoV-2 mutations altering regulatory properties: deciphering host’s and virus’s perspectives

**DOI:** 10.1101/2020.06.15.150482

**Authors:** Abul Bashar Mir Md. Khademul Islam, Md. Abdullah-Al-Kamran Khan

**Author notes:** **Correspondence:** Dr. Abul Bashar Mir Md. Khademul Islam, Associate Professor, Department of Genetic Engineering and Biotechnology, University of Dhaka, Dhaka 1000, Bangladesh.

## Abstract

Since the first recorded case of the SARS-CoV-2, it has acquired several mutations in its genome while spreading throughout the globe. However, apart from some changes in protein coding, functional importance of these mutations in disease pathophysiology are still largely unknown. In this study, we investigated the significance of these mutations both from the host’s and virus’s perspective by analyzing the host miRNA binding and virus’s internal ribosome entry site (IRES), respectively. Strikingly, we observed that due to the acquired mutations, host miRNAs bind differently compared to the reference; where few of the miRNAs lost and few gained the binding affinity for targeting the viral genome. Moreover, functional enrichment analysis suggests that targets of both of these gained and lost miRNAs might be involved in various host immune signaling pathways. Also, we sought to shed some insights on the impacts of mutations on the IRES structure of SARS-CoV-2. Remarkably, we detected that three particular mutations in the IRES can disrupt its secondary structure which can further make the virus less functional. These results could be valuable in exploring the functional importance of the mutations of SARS-CoV-2 and could provide novel insights into the differences observed different parts of the world.

## Introduction

The recent COVID-19 pandemic has posed a serious threat to the public health sector around the whole globe as it has already spread to 213 countries/territories. Approximately, 7.8 million people got infected with SARS-CoV-2 and half a million suffered death from this till the date of writing this article [1]. The woes of this pandemic are still on the rise as the detailed molecular aspects of the pathogenesis are still intangible.

Since the reporting of the first genome sequence of SARS-CoV-2, a huge number of genomes are being sequenced everyday. Comparing to the reference, genome sequences from different parts of the globe have suggested that this virus might have been acquiring many sequence alterations while spreading. But the functional significance of these variations are largely unknown and are not well-correlated with the disease pathobiology or infectivity.

MicroRNAs are small non-coding RNA (ncRNAs) which can play important roles during a viral infection. miRNAs can provide benefit for the host by exerting antiviral mechanisms through the innate and adaptive immune systems [2]; on the other hand, they can also facilitate the viral propagation by negatively modulating the host immune responses [3]. It was also reported that mutations in the miRNA binding sites in the viral genome can expedite the viral proliferation during an infection [2].

Ribosomal frame-shifting using the internal ribosome entry site (IRES) has previously been reported as important for coronaviruses as they use this site for synthesizing their proteins [4]. Moreover, alterations in the IRES can result in lethal effects on the SARS-CoV propagation and life-cycle [5]. Kelly et al. suggested that SARS-CoV-2 possess similar IRES structure like SARS-CoV [6].

In this present study, we sought to find out the comparative effects of the SARS-CoV-2 genome mutations on the basis of the host miRNA targeting profiles and IRES alterations. We illuminated the putative alterations of the host-miRNA binding profiles that might have arose due to the acquired mutations in different world-wide isolates. Also, the probable structural alterations due to the variations in SARS-CoV-2 IRES were investigated.

## Materials and Methods

### Mining of SNPs and their world-wide frequencies

We extracted the SNPs of SARS-CoV-2 which have world-wide frequency ≥ 10 for the miRNA binding analysis; while for analyzing the effects on IRES structure, we took all SNPs found in the IRES region of SARS-CoV-2. We obtained these information from “The 2019 Novel Coronavirus Resource” database along with the SNP annotations as on 30^th^ April, 2020 which included information from 8614 “High quality” (As set by GISAID database: Complete and high coverage sequences only with < 1% Ns, <0.05% unique amino acid mutations, and no insertion or deletion unless confirmed by submitter) SARS-CoV-2 genomes [7].

### Viral RNA-host miRNA interaction analysis

We have only taken fragmented viral sequences around our targeted viral reference sequence and SNPs (20 nts upstream and 20 nts downstream of a SNP) for the interaction analysis. Then we used three different tools, namely-RNAhybrid [8], miRanda [9], IntaRNA [10] for analyzing the miRNA-RNA interaction. We considered those as high confidence interaction when it has been predicted by every tools that we used and have values- (i) for RNAhybrid: MFE ≥ −35 KJ/mol and p-value < .001, (ii) for miRanda: energy ≥ −15 KJ/mol, (iii) for IntaRNA: ΔΔG ◻≤ −15 KJ/mol. We compiled the commonly predicted miRNAs from these three tools using the respective abovementioned cut-offs to reduce the false positives predictions.

### Retrieval of country-wise COVID-19 statistics

COVID-19 related statistics (Total deaths, Deaths/1 Million population, Case fatality rate) from different countries were extracted from the Worldometer website [1] on 24^th^ April, 2020. We presented the country-wise COVID-19 fatality statistics along with the miRNA binding profiles around the targeted SNPs of the associated country.

### Target genes functional enrichment analysis

We performed the functional enrichment of the experimentally validated targets (obtained from mirTarBase database [11]) of miRNAs that can bind around the SNPs of SARS-CoV-2 using Gitools v1.8.4 [12] utilizing KEGG and Bioplanet pathway modules. Resulting p-values were adjusted for multiple testing using the Benjamin and Hochberg’s method of False Discovery Rate (FDR).

### Prediction of the secondary structures of the IRES

The coordinates of the experimentally validated IRES of SARS-CoV-2 was extracted from previously conducted study [6] and adjusted accordingly. While predicting the secondary structures of the IRES incorporating the reference and the mutated bases, we took 50 additional nucleotides both upstream and downstream from the coordinates of the IRES. We used RNAfold tool [13] for the prediction of the IRES secondary structures.

## Results and Discussion

We have obtained 377 high confidence (mutation frequency ≥10) SNPs from the database and used the regions around those for miRNA binding analysis (Data not shown). Not all SNPs could alter the miRNA binding affinity. Ten miRNAs were observed to bind both the targeted reference and mutated regions of the SARS-CoV-2 genome (Supplementary file 1). Five miRNAs were found to bind only the regions of the reference genome which were not found significant for the mutated sequences, where 5 different other miRNAs can bind to the regions around the observed mutations but not found in the reference sequences (Figure 1). We have considered the miRNAs as “Gained” ones which can bind the mutated sequences; and miRNAs which were not found to bind the mutated sequences but can bind the reference are termed as “Lost” miRNAs. Most of the miRNAs binding were observed for the targeted regions of the N and ORF1ab gene sequence. Also, miRNAs were also found to bind the targeted regions of ORF3a, ORF7a, and 3’UTR (Figure 1).

**Figure 1:**
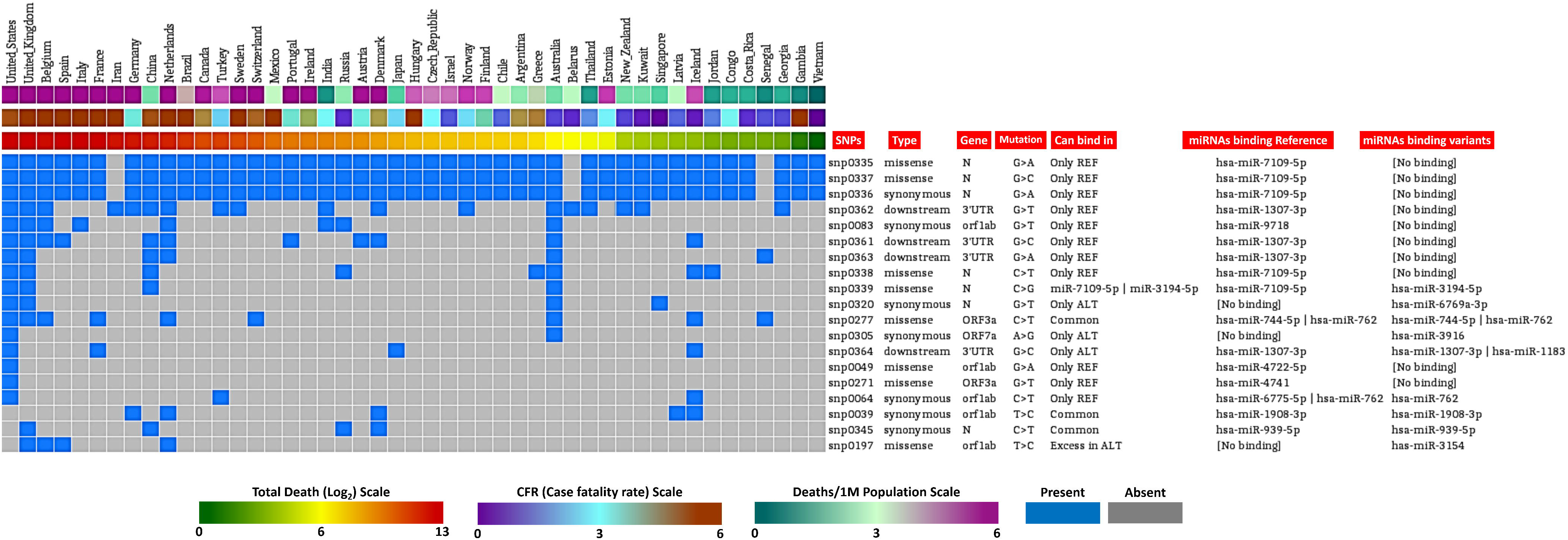
Country-wise distribution of the analyzed SNPs and the predicted miRNA bind profile around the regions of these SNPs. Country-wise deaths/1M population, CFR, and total deaths are also provided in color coded heatmaps.

We observed that the countries in which these mutations were much prevalent (Supplementary file 2), a less number of miRNAs can bind to the SARS-CoV-2 genome/transcripts due to the acquired mutations, whereas these miRNAs can bind to the associated reference sequences (Figure 1). It was also detected that countries having high fatalities from COVID-19 have more mutations, and maybe these mutations can resist the binding of host miRNAs and giving those viruses with the mutations a competitive edge over the reference virus. Still, more experimental evidences should be searched in order to find out the definitive roles of these mutations in escaping from the host antiviral miRNAs.

Moreover, the functional enrichment analysis using the targets of these found miRNAs suggested that due to the loss of miRNA binding can impact the host adaptive and innate immune signaling pathways, hypoxia response etc. (Figure 2A-B); whereas targets of the gained miRNAs for the mutations are also involved in pathways like-interleukin signaling, autophagy etc. (Figure 2A-B). So, the acquired mutations can lead to the alterations in miRNAs which can in turn result in the altered host immune responses during the viral infection.

**Figure 2:**
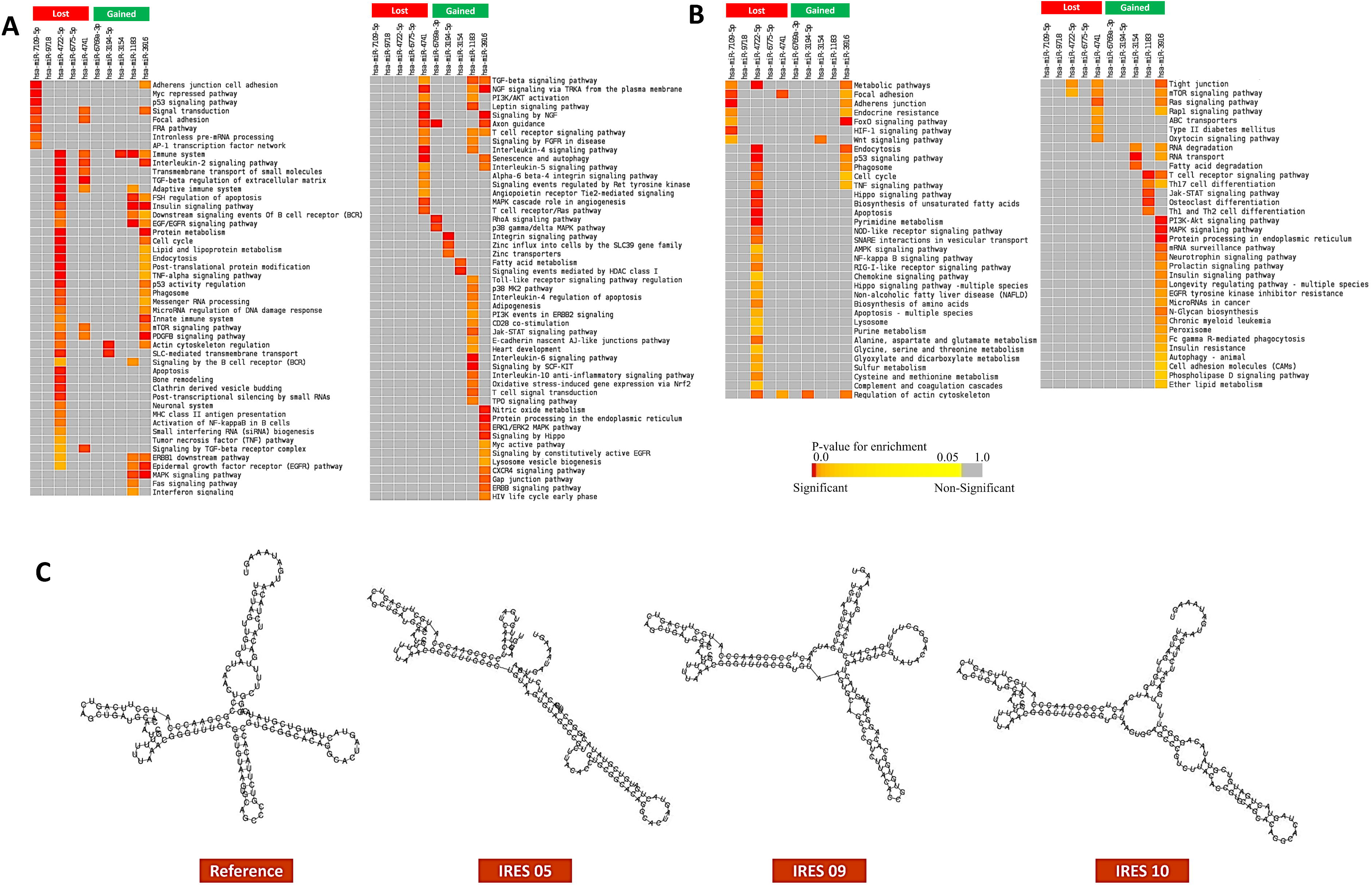
Functional enrichment analysis using the targets of host miRNA which can bind differentially around the mutated regions of SARS-CoV-2 utilizing **A.** KEGG pathway and, **B.** Bioplanet pathway modules. **C.** Secondary structures of the IRES of SARS-CoV-2 for the reference and mutated sequences. Significance of enrichment in terms of adjusted p-value (< 0.05) is represented in color coded P-value scale for all heatmaps. Color towards red indicates higher significance and color towards yellow indicates less significance, while grey means non-significant. Only selected significant enriched terms are shown.

We scrutinized 19 mutations in the IRES region of SARS-CoV-2 which was previously defined by Kelly et al. [6]. We aimed to illustrate the impact of the mutations in the secondary structure of the IRES of SARS-CoV-2 and predicted the secondary structures using RNAfold tool [13]. Interestingly, we observed that among the 19 mutations, 3 can significantly alter the secondary structure of the IRES of SARS-CoV-2 (Figure 2C). These 3 mutations in the IRES region are 13487: C>T, 13506: C>T, and 13507: G>A (Supplementary file 3). Mutations in these positions can destroy the viral translation process and render the virus less virulent. As, the studies regarding the IRES alterations are still in infancy, so more targeted researches should be conducted to elucidate the functional importance of these variations.

In our current analysis, we focused on the functional significance of the mutations reported for SARS-CoV-2 on the basis of host miRNA binding and the putative alterations in the IRES structure. Our preliminary results can suggest that these mutations can alter the binding pattern of host miRNAs which can ultimately result in alternative responses from these binding. Also, we revealed that several mutations in the IRES can disrupt its secondary structure which might suggest why the virus is affecting less in some countries compared to the others. Taking these observations in count, further studies should be conducted to completely understand the functional effects of the acquired mutations in the SARS-CoV-2 genome, and integration of other clinical information with these results can provide more insights on the dynamics of the SARS-CoV-2 infection.

## Supporting information

Ten miRNAs were observed to bind both the targeted reference and mutated regions of the SARS-CoV-2 genome

We observed that the countries in which these mutations were much prevalent

These 3 mutations in the IRES region are 13487: C>T, 13506: C>T, and 13507: G>A

## Author’s contribution

ABMMKI conceived the project, designed the workflow and performed the analyses. Both authors wrote the manuscript. All authors read and approved the final manuscript.

## Acknowledgments

We would also like to thank all the authors who have kindly deposited and shared genome data on GISAID (https://www.gisaid.org/).

## Data Availability Statement

Publicly available data were utilized. Analyses generated data are deposited as supplementary files.

## List of supplementary files

**Supplementary file 1:** SNPs, their frequencies and the miRNAs which can bind around these SNPs.

**Supplementary file 2:** Frequencies of the SNPs along with their associated countries’ fatality statistics.

**Supplementary file 3:** List of mutations found in the IRES of SARS-CoV-2 and their frequencies.

## References

1. Worldometer, Coronavirus Cases. 2020. p. 1–22.

2. Trobaugh, D.W. and W.B. Klimstra, MicroRNA Regulation of RNA Virus Replication and Pathogenesis. Trends in molecular medicine, 2017. 23(1): p. 80–93.

3. Głobińska, A., M. Pawełczyk, and M.L. Kowalski, MicroRNAs and the immune response to respiratory virus infections. Expert review of clinical immunology, 2014. 10(7): p. 963–971.

4. Dinman, J.D., Mechanisms and implications of programmed translational frameshifting. Wiley interdisciplinary reviews. RNA, 2012. 3(5): p. 661–673.

5. Plant, E.P., et al., Altering SARS coronavirus frameshift efficiency affects genomic and subgenomic RNA production. Viruses, 2013. 5(1): p. 279–294.

6. Kelly, J.A. and J.D. Dinman, Structural and functional conservation of the programmed-1 ribosomal frameshift signal of SARS-CoV-2. bioRxiv, 2020: p. 2020.03.13.991083.

7. Zhao, W.M., et al., The 2019 novel coronavirus resource. Yi Chuan, 2020. 42(2): p. 212–221.

8. Krüger, J. and M. Rehmsmeier, RNAhybrid: microRNA target prediction easy, fast and flexible. Nucleic Acids Research, 2006. 34(suppl_2): p. W451–W454.

9. Betel, D., et al., The microRNA.org resource: targets and expression. Nucleic Acids Research, 2008. 36(suppl_1): p. D149–D153.

10. Mann, M., P.R. Wright, and R. Backofen, IntaRNA 2.0: enhanced and customizable prediction of RNA-RNA interactions. Nucleic acids research, 2017. 45(W1): p. W435–W439.

11. Huang, H.-Y., et al., miRTarBase 2020: updates to the experimentally validated microRNA–target interaction database. Nucleic Acids Research, 2019. 48(D1): p. D148–D154.

12. Perez-Llamas, C. and N. Lopez-Bigas, Gitools: analysis and visualisation of genomic data using interactive heat-maps. PloS one, 2011. 6(5).

13. Gruber, A.R., et al., The Vienna RNA websuite. Nucleic acids research, 2008. 36(Web Server issue): p. W70–W74.

